# Human iPSC-derived brain endothelial microvessels in a multi-well format enable permeability screens of anti-inflammatory drugs

**DOI:** 10.1101/2021.05.03.442133

**Authors:** Sven Fengler, Birgit Kurkowsky, Sanjeev Kumar Kaushalya, Wera Roth, Eugenio Fava, Philip Denner

**Affiliations:** Laboratory Automation Technologies (LAT) – Core Research Facilities and Services (CRFS), German Center for Neurodegenerative Diseases (DZNE), 53127 Bonn, Germany; Light Microscopy Facility (LMF) – Core Research Facilities and Services (CRFS), German Center for Neurodegenerative Diseases (DZNE), 53127 Bonn, Germany

**Keywords:** Blood-brain barrier, iPSC, conditioned medium, organ-on-a-chip, microfluidic, high-content, drug permeability, neurodegenerative diseases, anti-inflammatory drugs

## Abstract

Optimizing drug candidates for blood-brain barrier (BBB) penetration in humans remains one of the key challenges and many devastating brain diseases including neurodegenerative diseases still do not have adequate treatments. So far, it has been difficult to establish state-of-the-art human stem cell derived *in vitro* models that mimic physiological barrier properties including a 3D microvasculature in a format that is scalable enough to screen drugs for BBB penetration in early drug development phases. To address this challenge, we established human induced pluripotent stem cell (iPSC)-derived brain endothelial microvessels in a standardized and scalable multi-well plate format. iPSC-derived brain microvascular endothelial cells (BMECs) were supplemented with primary cell conditioned media and grew to intact microvessels in 10 days of culturing. Produced microvessels show a typical BBB phenotype including endothelial protein expression, tight-junctions and polarized localization of efflux transporter. Microvessels exhibited physiological relevant trans-endothelial electrical resistance (TEER), were leak-tight for 10 kDa dextran-Alexa 647 and strongly limited the permeability of sodium fluorescein (NaF). Permeability tests with reference compounds confirmed the suitability of our model as platform to identify potential BBB penetrating anti-inflammatory drugs. In summary, the here presented brain microvessel platform recapitulates physiological properties and allows rapid screening of BBB permeable anti-inflammatory compounds that has been suggested as promising substances to cure so far untreatable neurodegenerative diseases.

## Introduction

The BBB strictly regulates the molecular traffic between the blood and the brain and ensures a homeostatic environment that is essential for healthy brain function. The BBB consists of tightly connected brain-specific endothelial cells which seal the brain vasculature. BMECs express unique tight-junction proteins limiting paracellular diffusion, nutrient transporter regulating the flow of energy and efflux transporter preventing entry of unwanted toxic substances from the brain (Abbott et al., 2010; Pardridge, 2005). While this barrier is essential for preserving healthy brain activity, it has been estimated that only 2 % of small molecule drugs can cross the human BBB and reach a therapeutic target (Pardridge, 2001, 2020). So far, many devastating brain diseases including neurodegenerative diseases do not have adequate treatments except for some symptomatic therapies (Bright et al., 2019; Dong et al., 2019; Pangalos et al., 2007). More and more experimental evidence supports the concept that neuro-inflammation is involved and connected to the disease progression of neurodegeneration (Bright et al., 2019; Heneka et al., 2015; Venegas et al., 2017). This collective knowledge suggests that anti-inflammatory drugs could potentially treat neurodegenerative diseases (Voet et al., 2019). Thus, the development of therapeutic approaches that specifically target inflammatory signaling in the brain is now in focus of intense biological and clinical interest (Kudelova et al., 2015; Vila and Przedborski, 2003). However, the lead optimization for brain penetration remains one of the major challenges in drug discovery. The common prediction models for brain penetration is based on an integration of several *in vivo* and *in vitro* parameters including unbound drug concentration, intra-brain distribution and BBB permeability (Reichel, 2009), whereas the latter property remains the principal prerequisite for every new central nervous system (CNS) drug. Thus, the development of advanced *in vitro* models to screen compounds for BBB permeability are in focus to finally optimize and better predict brain penetration of drugs in humans. Moreover, drug discovery for non-CNS drugs would benefit from same *in vitro* models to finally screen for substances that cannot cross the human BBB after systemic application.

Various BBB models based on primary brain endothelial cells from animals have contributed to the development of drug permeability assays (Deli et al., 2005), but due to their species-specific differences in brain physiology human models should be favored in drug discovery (Uchida et al., 2011). However, human models with primary BMECs are also not ideal because these cells are limited in scale due to their availability of biopsied brain tissue (Bernas et al., 2010). Moreover, models with immortalized primary BMECs are scalable, but generally provide only a suboptimal barrier tightness indicated by a low TEER (Weksler et al., 2005). In contrast, TEER values in a range of 1000-6000 Ωxcm^2^ were measured in rodent brain vessels, highlighting a significant discrepancy with most *in vitro* models (Butt et al., 1990).

Apparently, current technologies are not sufficient to fully reproduce the complex physiology of the human BBB. However, interdisciplinary achievements in biotechnology, materials engineering and availability of new human cell sources have enabled the development of innovative and highly integrated almost physiological *in vitro* BBB models recently (Kaisar et al., 2017). During the last years, BMECs differentiated from various human stem cell sources have been used for *in vitro* models with promising BBB properties (Boyer-Di Ponio et al., 2014; Cecchelli et al., 2014; Katt et al., 2016; Lippmann et al., 2012). In particular, human iPSC-derived BMEC co-cultures combined with additional human cell types such as pericytes and astrocytes indicated physiological barrier properties and tightness as shown in rodent models (Lippmann et al., 2014). One important advantage of stem cell-derived models is that they originate from renewable cultures and therefore overcome material challenges required for upscaling.

Most human BBB *in vitro* models and also those using recent iPSC technologies are based on static cultures in Transwell plates with a 2D endothelial layer that grows on an artificial filter membrane to separate an apical from a basolateral side. However, microvascular endothelial cells *in vivo* are continuously exposed to wall shear stress, a tangential force generated by the flow of blood. It has been reported that applied shear stress in culture positively influenced the barrier tightness of immortalized murine or human brain endothelial cells growing in microfluidic devices (Booth and Kim, 2012; Griep et al., 2013). Moreover, perfused iPSC-derived BMEC microvessels in hydrogel scaffolds showed a significant lower permeability for applied tracer molecules compared to static cultures (Faley et al., 2019). Efforts to simulate more realistic representation of the BBB morphology by reproducing the microcirculatory environment of the brain to account for liquid flow and shear stress have led to the development of advanced microfluidic systems with various cell sources including human primary cells combined with iPSC-technology (Brown et al., 2020; Grifno et al., 2019; Lee et al., 2020; Park et al., 2019; Vatine et al., 2019). The main advantage of these microfluidic chambers is that they are addressable by common microscopy techniques and applications range from high resolution tissue characterization to quality control assays with fluorescent tracer molecules. However, microfluidic systems are generally limited in terms of throughput due to their complex setup making them relatively unsuitable for screening purposes. Recent microplate development overcomes this limitation by providing perfused microfluidic chambers in a standardized multi-well plate format (Koo et al., 2018; Wevers et al., 2018).

In the following work, we present the development of perfused human iPSC-derived brain endothelial microvessels in microfluidic chambers arranged in a 384-well plate format to screen drugs for potential BBB permeability. Our protocol allows the parallel production of 40 microvessels per plate within 10 days that includes a maturation step with standardized human brain vascular pericyte (HBVP) conditioned media. Microvessels revealed an endothelial protein expression, tight-junctions as well as polarized localization of efflux transporter in absence of complex primary cell co-cultures. Microvessels were leak-tight for 10 kDa dextran conjugated to Alexa 647 and strongly limited the permeability of NaF. TEER of microvessels obtained physiological relevant values that correlated with NaF data. In conclusion, the here presented human iPSC-derived brain microvessel platform recapitulated physiological properties and is suitable to screen potential anti-inflammatory compounds for BBB permeability in a standardized microtiter plate format.

## Results

### BMEC maturation by conditioned media results in similar barrier tightness compared to co-cultures

To develop functional BBB endothelial cells from human stem cells, we differentiated IMR90-4 iPSCs to BMECs in a four-step process by following the protocol published by *Stebbins et al*., *2016* (Supplementary Figure S1A). After 3 days (D3) of iPSCs expansion, a heterogenous cell culture containing endothelial cell and neuronal cell progenitors was generated by applying unconditioned medium (UM) for 6 days (Supplementary Figure S1B, Supplementary Video V1). Endothelial cells were selectively expanded by adding endothelial cell medium supplemented with all-trans retinoic acid (RA) and bFGF (EC+/+). Cultures were finally subcultured on collagen/fibronectin on day 8 (D8) as previously described (Stebbins et al., 2016). To characterize the resulting BMEC phenotype after differentiation, cells were stained for BBB marker proteins. We found a positive stain for typical endothelial markers (PECAM-1 and Ve-Cadherin), tight-junction proteins (Occludin, ZO-1 and Claudin-5), nutrient transporter (Glut-1) and efflux transporters (P-gp and BCRP) (Figure 1A). To investigate the optimal time point of barrier tightness after subculturing, TEER of BMECs growing apical on 2D Transwell-filter was recorded from day 9 to day 12 (D9 - D12). Average values of approximately 2500 Ωxcm^2^ were obtained at day 10 (D10) (Figure 1B). Treatment with Staurosporine (STS) resulted in disruption of tight-junctions that was detectable by imaging and the loss of TEER (Supplementary Figure S1C). Overall, our results indicated that iPSC-derived BMECs express relevant BBB endothelial marker and reveal physiological barrier properties at D10.

**Figure 1.**
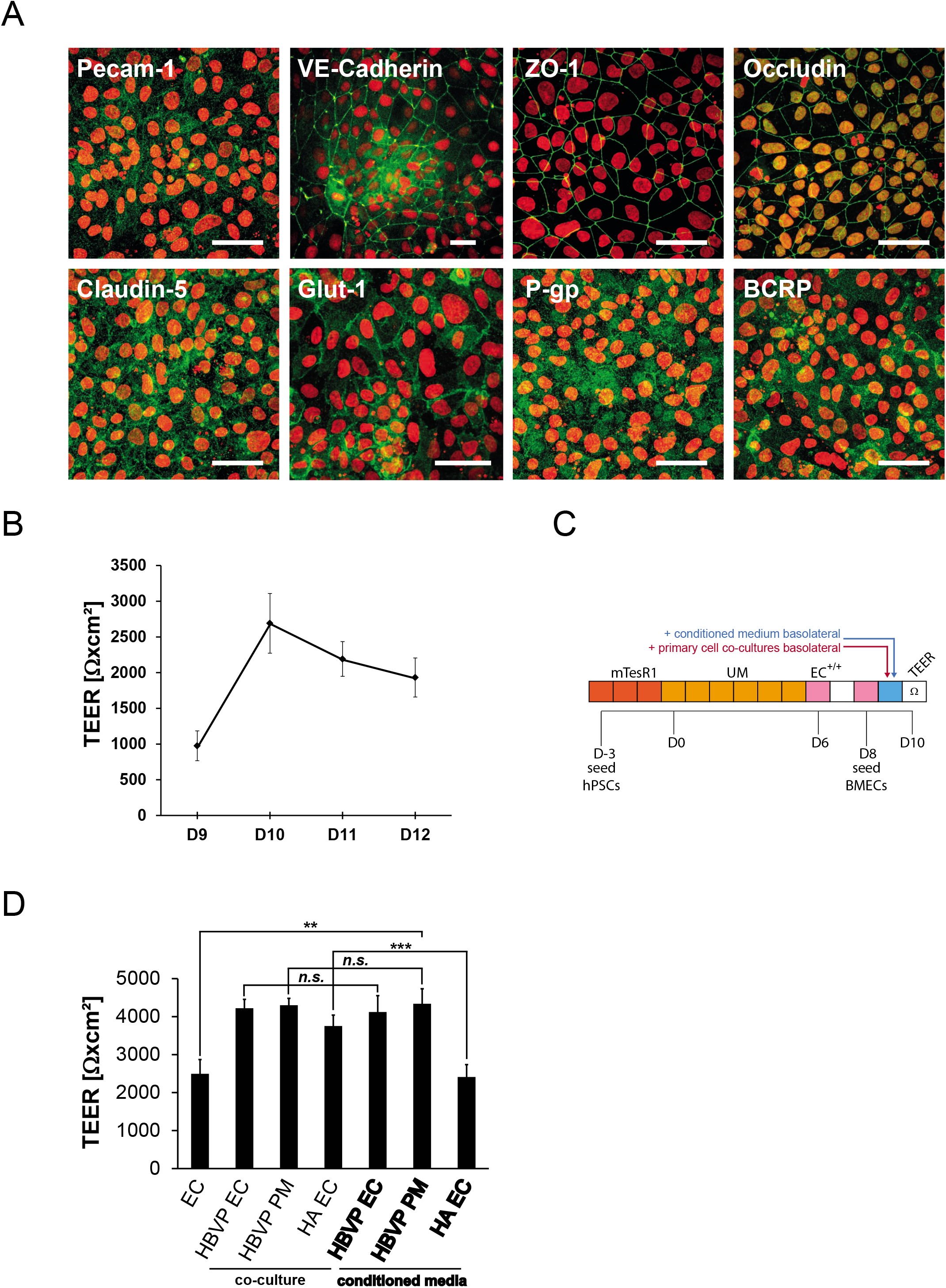
Characterization of BMEC cultures and maturation with primary cell conditioned media. (A) Immunocytochemistry of BMEC monolayer cultures at D10. Staining’s of endothelial marker (PECAM-1, VE-Cadherin); tight-junction and associated proteins (ZO-1, Occludin, Claudin-5); nutrient transporter (Glut1) and efflux transporter (P-gp, BCRP) are shown. Scale bar equals to 50 µm. (B) TEER of BMEC cultures on Transwell chambers from D9 to D12. Filter duplicates were measured and averages were calculated. Error bars indicate the standard deviation over three independent experiments (n=3). (C) Modified differentiation protocol including the maturation step with either primary cell conditioned media or classical co-culture on basolateral chambers. (D) Comparison of TEER when BMEC cultures were matured in co-cultures or with primary cell conditioned media of either HBVP or HA. Duplicate filters were used to calculate averages. Error bars indicate the standard deviation of three independent experiments (n=3). Statistical significance was calculated using the unpaired T-test (** p<0.005; *** p<0.0005; n.s. p>0.05). Abbreviations: HBVP: human brain vascular pericytes, HA: human astrocytes, EC: Endothelial cell medium, PM: Pericyte Medium).

Recent, iPSC-derived BMEC co-culture systems with additional cell types of the microvascular environment like pericytes or astrocytes have reached TEER values that were comparable to those measured in rodent brain vessels (Butt et al., 1990; Lippmann et al., 2014). However, human primary brain cells in a co-culture system introduce an additional level of complexity to the experiment setup. Moreover, compound effects measured in a heterotypical cell mode are often very difficult for data interpretation. To overcome such limitations, we investigated whether primary cell conditioned media derived from human brain vascular pericytes (HBVP) and human astrocytes (HA) could provide similar TEER as shown in classical co-cultures. A detailed protocol for the production of conditioned medium is depicted in the methods section. In brief, either 4×10^6^ HBVPs or 1×10^6^ HAs were cultured with endothelial cell medium without RA and bFGF (EC−/−). The applied EC−/− medium was exchanged after culturing for 2 days and first the conditioned medium was harvested on day 3 after seeding. The harvested conditioned media was always stored at −20°C. For the experiment, Transwell filters with apical BMEC cultures were either supplemented with conditioned medium or filters were placed on top of primary cells growing in basolateral chambers (Figure 1C). We found, that co-cultures with HBVPs or HAs significantly increased TEER by 1.7 and 1.5 fold, respectively when compared to conventional BMEC cultures without a co-culture (Figure 1D). Furthermore, BMEC barriers supplemented with HBVP conditioned EC medium showed significantly enhanced TEER values >4000 Ωxcm^2^, similar to HBVP co-cultures. Same results were obtained when ScienCell™ provided pericyte medium (PM) was previously conditioned with HBVPs and applied. Interestingly, conditioned medium from HA had no supporting effect compared to HA co-cultures, demonstrating different supporting mechanisms. In summary, these data indicated that HBVP conditioned media enhances BMEC barrier tightness significantly and to the same extent as shown for classical co-cultures. To investigate whether HBVP conditioned medium also provides reliable permeability values of BMEC cultures on Transwells, we next determined the permeability (PE) of the commonly used fluorescent tracer NaF (Kaya and Ahishali, 2011). NaF was supplemented on apical chambers and medium from basolateral chambers was collected every 15 min over a period of 1 hr. The fluorescence intensity of basolateral media was measured and the PE was calculated as described in the methods section. We found no significant differences in NaF permeability when BMECs were either supplemented with HBVP conditioned medium or co-cultured with HBVPs (Supplementary Figure S1D). Both culture systems reached average PE values between 4.4 – 6.5 x 10^−7^ cm/s representing typical values for iPSC-derived BMEC Transwell-cultures (Stebbins et al., 2016). These data collectively indicated that functional BMEC barriers can be derived from iPSCs and require 10 days to fully mature. Furthermore, barriers supplemented with HBVP conditioned media showed physiological TEER values like more complex co-cultures systems. In conclusion, our BMEC maturation protocol is favored for upscaling and standardizing screening applications because it circumvents the use of elaborate primary co-cultures in the assay.

### Perfused human iPSC-derived brain endothelial microvessels provide *in vivo* BBB characteristics

We demonstrated that our differentiated iPSC-derived BMEC monolayer provide a functional barrier in 2D Transwell systems. However, it has been reported that applied shear stress positively influences BBB properties in 3D vessel-like endothelial cultures (Faley et al., 2019; Griep et al., 2013). Moreover, a perfused BBB *in vitro* model, suitable for high-content applications has been previously reported with immortalized human brain endothelial cells (Wevers et al., 2018). Therefore, we next transformed our 2D permeability model to a miniaturized screening format based on perfused 3D brain endothelial microvessels to mimic *in vivo*-like BBB characteristics. For the production of iPSC-derived BMEC microvessels, three-lane OrganoPlates® (Mimetas) were used that allow a parallel production of 40 microvessels per plate. A detailed description of the established procedure can be found in the methods section. In brief, gel channels were filled with collagen extracellular matrix (ECM) and differentiated BMECs were seeded at D8 into top channels (Figure 2A). After 24 hrs culturing in EC+/+, medium was exchanged by adding EC−/− supplemented with HBVP-conditioned PM (1:4 ratio). We used HBVP conditioned PM for microvessel maturation since more reproducible TEER values were measured in Transwell experiments (data not shown).

**Figure 2.**
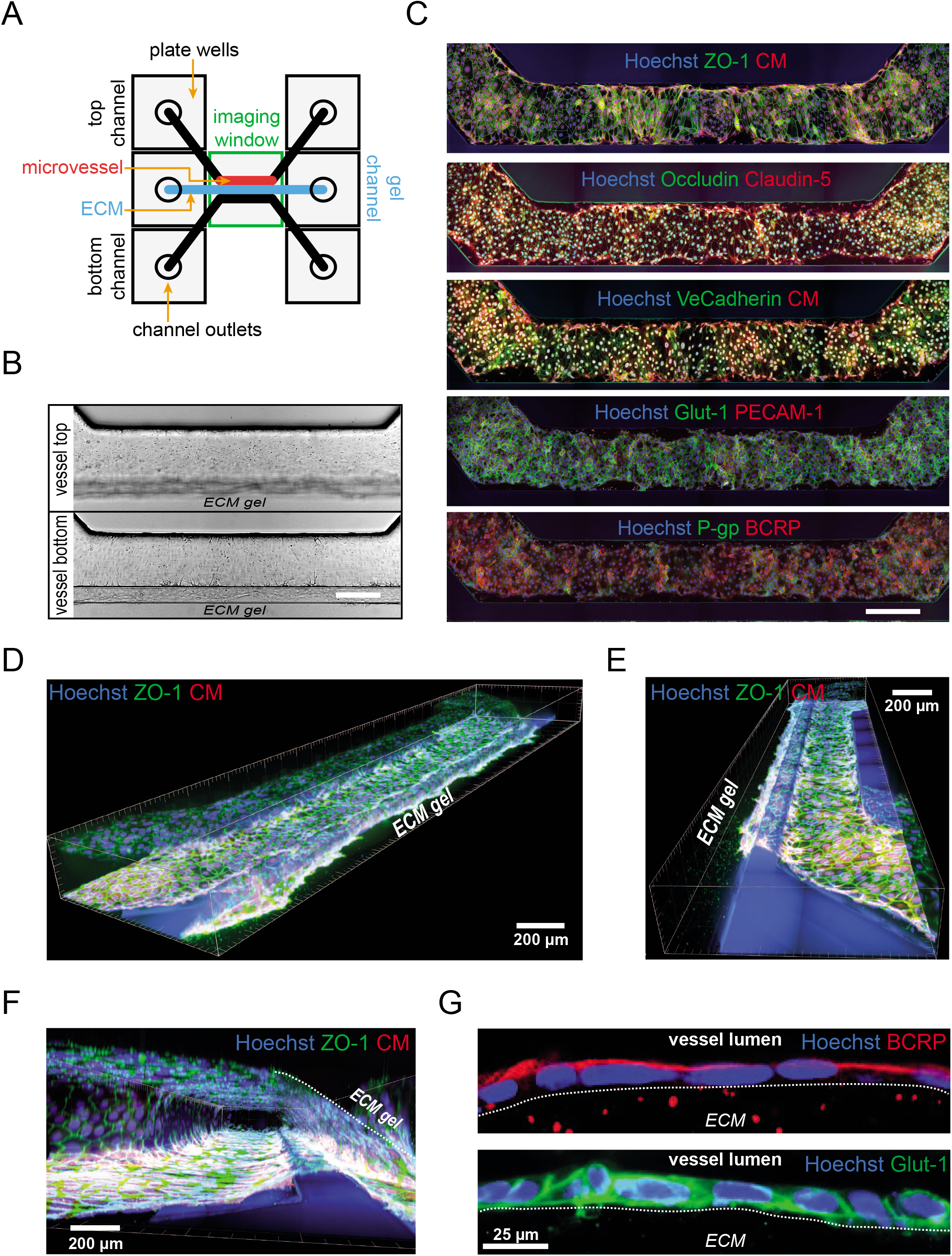
Characterization of perfused human iPSC-derived brain endothelial microvessels. (A) Schemata of a single three-lane OrganoPlate® chamber containing a microvessel (red) inside the top channel. The microvessel grows in contact to the ECM gel channel (light blue). (B) Bright-field images of microvessels, 48 hrs after cell seeding. Representative images at 20 µm (vessel bottom) and 170 µm (vessel top) height from channel bottom are shown. Scalebar equals to 300 µm. (C) Immunocytochemistry of microvessels 48 hrs after cell seeding. Representative images (from the microvessel bottom) of tight-junction proteins (ZO-1, Occludin, Claudin-5), endothelial marker (VE-Cadherin, PECAM-1), nutrient transporter (Glut-1) and efflux transporter (P-gp, BCRP) expression are shown. Nuclei staining with Hoechst and cytoplasm staining with CellMask™ as indicated. Scalebar equals to 300 µm. (D-F) 3D imaging of microvessels stained with nuclei (Hoechst), tight-junction (ZO-1) and cytoplasm CellMask™ marker. (G) Subcellular localization of BCRP and Glut-1. Microvessel wall sections at the vessel/ECM contact are shown.

Next, microfluidic plates were placed on a perfusion rocker (Mimetas) inside the incubator to start medium perfusion within the plate channels. After 48 hrs (D10), closed microvessels were observed by microscopy (Figure 2B). To investigate whether produced microvessels consist of cells with a functional BMEC phenotype, microvessels were stained by immunofluorescence (Figure 2C). 3D imaging demonstrated the expression of endothelial marker (PECAM-1 and Ve-Cadherin), tight-junction proteins (Occludin, ZO-1 and Claudin-5), nutrient transporter (Glut-1) and efflux transporters (P-gp and BCRP). This result indicated that vessel-cells express relevant BBB endothelial proteins. To demonstrate whether the entire surface of perfused microvessels and in particular the area connected to the ECM was closed by a BMEC layer, 3D image reconstruction was performed. We found that the entire microvessel surface was closed by a BMEC-monolayer in which single cells were connected by tight-junction proteins (Figure 2D-F, Supplementary Video V2). The microvessel covered the entire top channel lumen including the ECM contact side. Next, the subcellular localization of efflux transporter (Breast Cancer Resistant Protein) BCRP and nutrient transporter Glut-1 was further inspected to investigate whether BMECs in microvessels have a polarized endothelial phenotype. It has been reported that BCRP is only localized at the luminal (blood-facing) plasma membrane in contrast to Glut-1 which is present at the luminal but also at the abluminal (brain-facing) plasma membrane (Iorio et al., 2016; Simpson et al., 2007). Consistent with previous observations *in vivo*, we observed a specific luminal localization of BCRP and a luminal-abluminal localization of Glut-1 (Figure 2E). These results indicated a polarization of BMECs within perfused microvessel cultures. Overall, our data collectively demonstrated that seeded iPSC-derived BMECs matured to functional and polarized microvessels with a monolayer thickness held together by tight-junction proteins.

### Precise determination of microvessel tightness by fluorescent tracer and TEER reveals *in vivo*-like barrier properties

After characterization of microvessels, we next investigated the vessel functionality by addressing its integrity which is essential for drug permeability studies. One of the major advantages of using microfluidic plates is that vessel integrity can be analyzed by imaging of inert fluorescent tracer. As tracers, we choose 10 kDa dextran conjugated to Alexa 647 (DEX-A647) and the small 376 Da NaF. Both tracers have been described as only poorly BBB penetrating in rodents, whereas the permeability of 10 kDa dextran is more compromised in comparison to NaF (Shi et al., 2014; Yen et al., 2013; Yuan et al., 2009). For integrity tests of microvessels, growth medium was supplemented with both tracers and added into top wells on D10. Plates were further incubated under perfusion condition and both tracer intensities were imaged after 30, 150 and 300 min (Figure 3A). Intensity values from the top and the gel chamber were measured by automated image analysis as described in the methods section. Tracer mean intensity ratios [gel/top] were calculated for each time point. The majority of microvessels maintained a low (<0.01) and constant DEX-A647 intensity ratio over the total assay duration of 300 min (Figure 3B and Supplementary Figure S2A). Such vessels were considered as DEX-A647 leak-tight. In contrast, vessels with higher or increasing intensity ratios over time were considered as DEX-A647 leaky. A subset of microvessels was treated with STS resulting in severe vessel damage and detachment from the channel surface (Supplementary Figure S2B). These STS treated microvessels showed an accelerated increase of DEX-A647 ratios (Figure 3 B). In parallel to DEX-A647, NaF intensity ratios were determined in the same set of micovessels to investigate NaF permeability in preselected DEX-A647 leak-tight and leaky vessels (Figure 3C and Supplementary Figure S2C). Previously considered DEX-A647 leaky-tight micovessels showed a slow linear increase (slope, *m = 0*.*004*) of NaF intensity ratio over time demonstrating very low penetrating properties. In summary, our data demonstrates that microvessels were leak-tight for 10 kDa DEX-A647 over several hours. Moreover, preselected DEX-A647 leak-tight microvessels strongly limited NaF penetration. Finally, the combination of both tracers facilitates the monitoring of the microvessel integrity over time.

**Figure 3.**
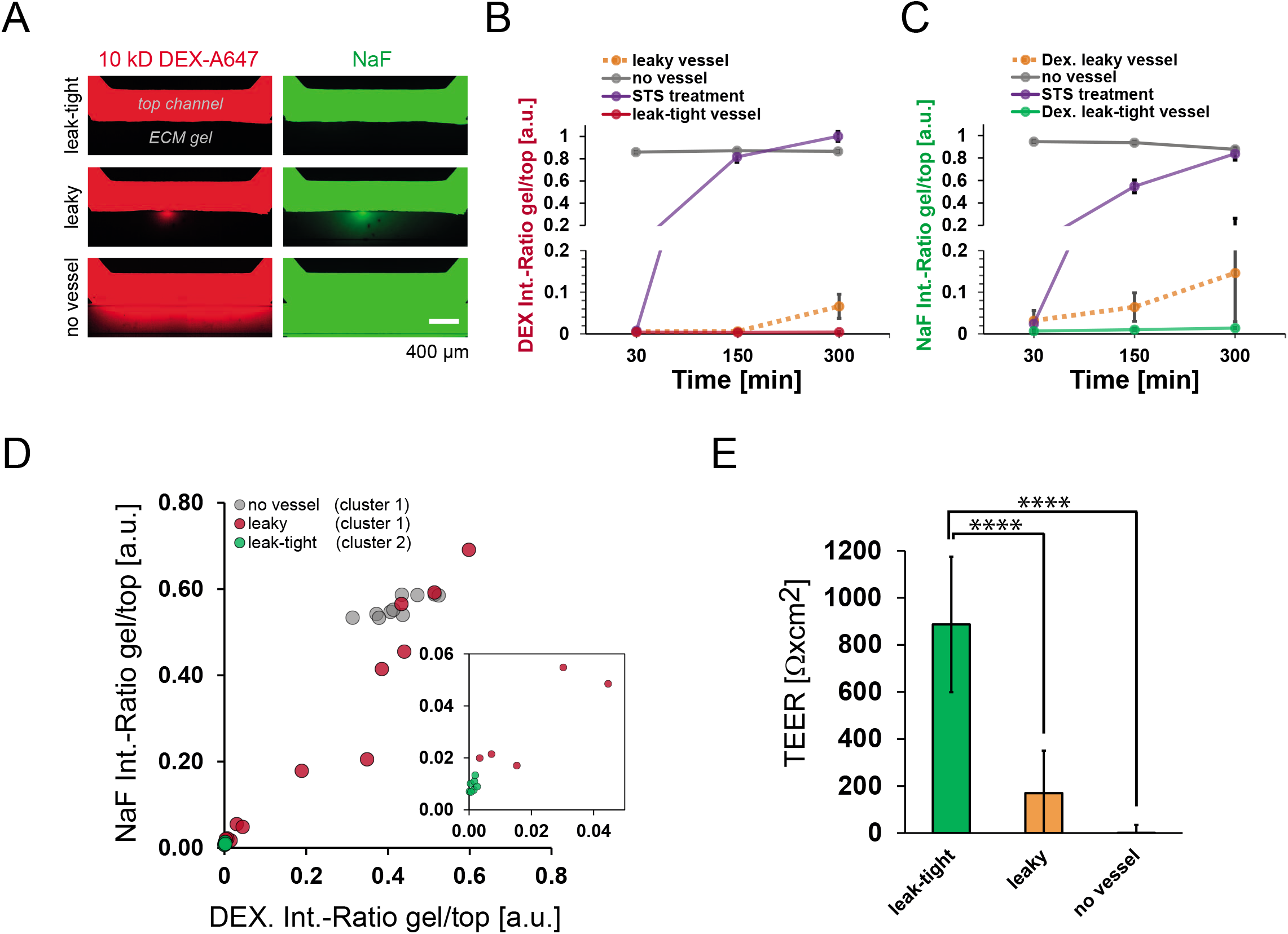
Perfused human brain endothelial microvessels reveals physiological barrier properties. (A) Representative images of a leak-tight and a leaky microvessel perfused with DEX-A647 and NaF in comparison to a chamber without microvessel (no vessel). (B) DEX-A647 mean intensity ratios (gel/top) over 300 min tracer incubation under perfusion. Dextran leak-tight vessels show a constant intensity ratio ~ 0.01 over time (red line). Leaky or STS treated vessels show ratios (> 0.01) that increase over time. Error bars indicate the SD of leak-tight vessels (n=6), leaky vessels (n=3), STS treated vessels (n=3) and chambers without microvessel (n=3). (C) NaF mean intensity ratios (gel/top) of previously identified DEX-A647 leak-tight and leaky vessels as shown in (B). NaF intensity ratios of DEX-A647 leak-tight vessels increased slowly over time. In comparison, NaF ratios of DEX-A647 leaky or STS treated vessels increased to values > 0.02. (D) K-mean clustering analysis to distinguish between leaky and leak-tight microvessels by using fluorescent tracer data (NaF and DEX-A647). Resulting clusters are represented in a 2D scatterplot of NaF versus DEX-A647 ratios after 300 min incubation. Cluster 1 contains all excluded leaky vessels (red) and also control chambers without microvessel (grey). Green dots indicate leak-tight members from cluster 2 after applying stringent thresholds according to incubation times. Smaller window represents the same dataset on a smaller scale. A total of n=26 microvessels were used for cluster and n=10 chambers without microvessel (no vessels). (E) TEER determined for leak-tight and leaky vessels in comparison to chambers without microvessels (no vessel) as indicated. Error bars indicate the SD of n=13 leak-tight, n=12 leaky and n=10 for no vessel chambers, respectively. Statistical significance was calculated by using the unpaired T-test (**** p<0.00001).

Previously considered DEX-A647 leak-tight microvessels had a DEX-A647 and NaF ratio < 0.02 and we observed a rather low variance of the ratio data as a common feature in this group. In contrast, DEX-A647 leaky or STS treated microvessels showed higher tracer ratios in general with a high variance between different microvessels that mirrors the broad range of vessel disruption. By considering the fact that all leak-tight microvessels have a distinct and rather narrow range for DEX-A647 and NaF ratios suggests that leak-tight vessels could be identified by using conventional cluster analysis. To determine DEX-A647 and NaF threshold values for leak-tight microvessels, a K-means clustering analysis was performed by combining 6 independent gel/top intensity ratio variants extracted from DEX-A647 and NaF images after two different incubation times (150 and 300 min). A detailed description of all ratio variants used in the cluster analysis can be found in the methods section. Finally, a set of 26 microvessels including preselected DEX-A647 leaky (R > 0.01) and leak-tight (R < 0.01) microvessels as well as 10 control chambers without a microvessel (no vessel) were used for clustering (Figure 3D). According to the clustering result and one-to-one image inspections stringent tracer ratio (R) thresholds to identify leak-tight microvessels were defined for different incubation times (R_gel/top_ DEX_150min_ < 0.01, R_gel/top_ DEX_300min_ < 0.01, R_gel/top_ NaF_150min_ < 0.015, R_gel/top_ NaF_300min_ < 0.02). By applying these stringent rules, we also excluded those cluster members found on the leak-tight-cluster boundaries (Supplementary Figure S2D). In summary, the presented clustering analysis was applied to determine reliable thresholds for both applied fluorescent tracer dyes to identify leak-tight microvessels in a quantitative and unbiased way. Furthermore, the determined thresholds provide a robust method to quality control microvessels for drug permeability tests.

To further evaluate the functionality of microvessels in real-time and tracer independent, TEER was assessed using a customized TEER measurement device. Detailed information about the device construction and measurement setup can be found in the methods section. In brief, the measurement was performed on a potential divider based resistance device. TEER was measured across the microvessel surface at the ECM gel contact by applying an Ag/AgCl-electrode into the right-top well and a second Ag/AgCl-electrode into the right-bottom well of each chamber (Supplementary Figure S2E). Finally, TEER was determined in leak-tight and leaky microvessels previously identified by cluster analysis. Average TEER values around 890 Ωxcm^2^ were measured for leak-tight microvessels and values around 170 Ωxcm^2^ for leaky vessels highlighting a significant difference between these two cases (Figure 3E). To demonstrate the sensitivity of the engineered TEER device, we correlated the NaF intensity ratios with corresponding TEER values of leak-tight microvessels (Supplementary Figure S2H). We obtained a linear correlation between NaF ratios and corresponding TEER values. This result could show that our TEER device provides sensitive measurements that enable the detection of small changes in microvessel tightness. For instance, leak-tight microvessels with lower NaF ratios (NaF_150min_ < 0.005 and NaF_300min_ < 0.01) reached TEER values > 1000 Ωxcm^2^ (Supplementary Figure S2I), indicating that these microvessels obtained physiological relevant TEER values (Butt et al., 1990; Mantle et al., 2016). For further verification of our TEER measurements, we perturbed a subset of leak-tight microvessels with STS. We found that, previously defined leak-tight microvessels shifted into the cluster of leaky vessels after STS treatment by considering NaF and DEX-A647 ratios from image data (Supplementary Figure S2J). In line with this observation, average TEER values of approx. 750 Ωxcm^2^ measured before treatment dropped to approx. 250 Ωxcm^2^ after treatment with STS. In summary, we could demonstrate that microvessels reached physiological relevant TEER values. TEER results correlated with NaF ratio data and allowed the detection of small changes in microvessel quality.

### Human iPSC-derived brain microvessels as model to determine drug permeability in a standardized high-content format

To demonstrate that produced microvessels were usable to screen for drug permeability, a collection of anti-inflammatory compounds targeting caspase-1, pan-caspase and the NLRP3 inflammasome were tested for microvessel permeation. As known BBB permeable reference, the NLRP3 inflammasome inhibitor CRID3 (Chen et al., 2017) was used. Furthermore, a set of three BBB impermeable peptide caspase inhibitors (Z-DEVD-FMK, Z-VAD(OH)-FMK and Ac-YVAD-CMK) were included. NaF, DEX-A647 and 40 µM compound were applied to the top wells of each chamber containing a matured microvessel (at D10). Microvessels were incubated for a total duration of 300 min under perfusion, 37°C and 5% CO_2_. NaF and DEX-A647 tracer intensities were imaged after 60, 150 and 300 min incubation. Tracer intensity ratios were determined to guarantee microvessel integrity by applying previously determined ratio thresholds. Tracer intensity ratios of microvessels containing compounds were within the previously defined cluster boundaries and were considered as leak-tight (Supplementary Figure S3A). After a total assay duration of 300 min, supernatants from basolateral bottom wells (receiver) were collected. We used a functional cytokine assay based on LPS-primed and activated human peripheral blood mononuclear cells (PBMCs) to identify anti-inflammatory compounds in the basolateral supernatants after a successful permeation. In brief, LPS in conjunction with the inflammasome activator ATP stimulates inflammasome mediated caspase-1 activity resulting in cleavage of IL-1β-precursor protein to its active and released form (Mangan et al., 2018). In case basolateral supernatants contained an anti-inflammatory compound, the reduction of IL-1β release was detected by homogeneous time-resolved fluorescence (HTRF). The potency of selected inflammasome inhibitors were verified in dose response in primed and activated PBMC before the actual permeability assay (Figure 4A). All compounds reduced the amount of IL-1β with IC:50 values in a range of 0.28 – 2.06 µM. To validate the IL-1β detection, the unperturbed PBMC IL-1β release was determined in presents of a neutralizing anti-IL-1β antibody. The level of unbound IL-1β correlated negatively to the titrated antibody concentration (Supplementary Figure S3B). Finally, the collected basolateral supernatants from permeability tests were applied on LPS-primed and activated PBMCs. Next, the concentration of released IL-1β was quantified to identify permeable compounds. The basolateral supernatants containing CRID3 significantly reduced the IL-1β release of PBMCs indicating its microvessel permeation, as expected (Figure 4B). On the other hand, all three impermeable reference peptide inhibitors (Z-DEVD-FMK, Z-VAD(OH)-FMK and Ac-YVAD-CMK) showed no permeability through microvessels. Furthermore, we identified impermeable properties for the broad spectrum peptidomimetic caspase inhibitor Emricasan. But surprisingly, two so far unknown anti-inflammatory compounds from our internal compound library showed a permeable profile (cmpd-A and cmpd-B). In summary, our data highlighted that the here presented human iPSC-derived brain microvessel platform is suitable as a tool to screen anti-inflammatory compounds for potential human BBB permeability.

**Figure 4.**
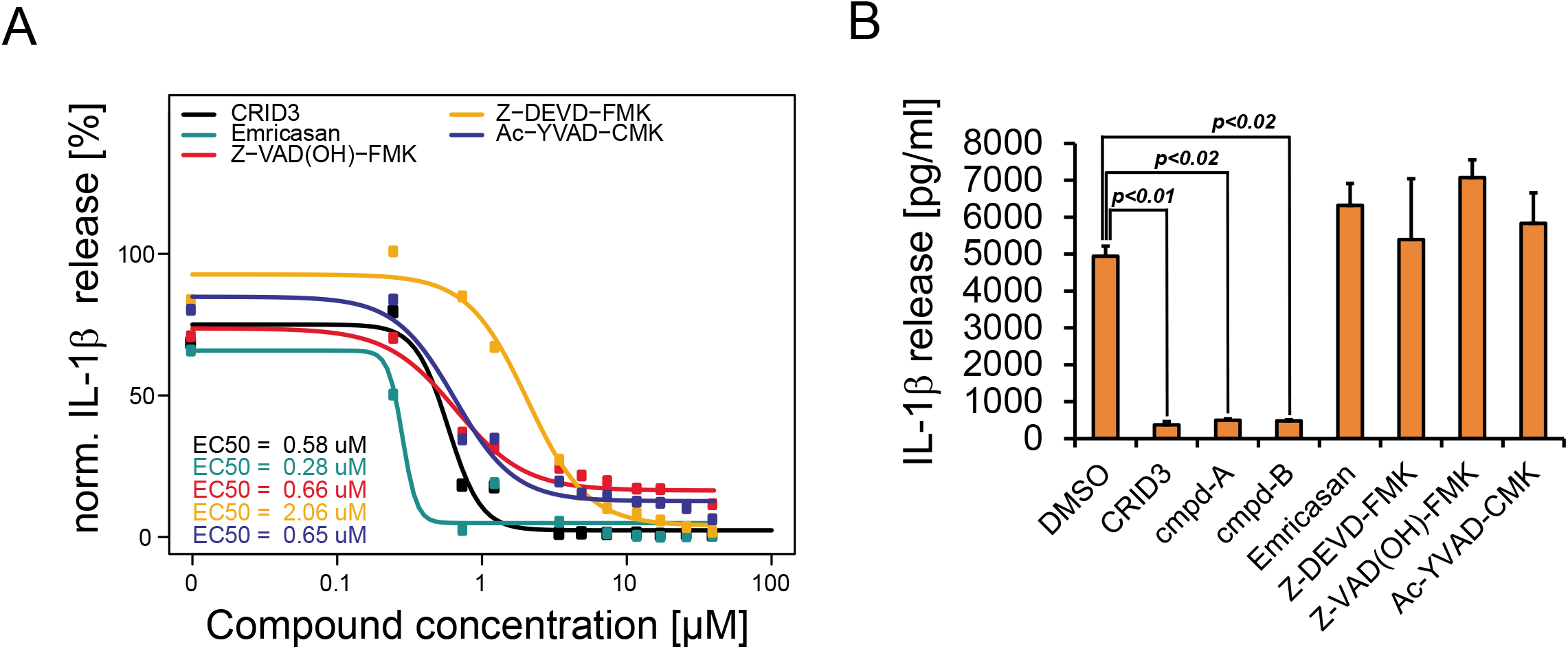
iPSC-derived microvessels as an *in vitro* model to identify BBB permeable anti-inflammatory compounds. (A) Dose response curves of selected anti-inflammatory compounds. IL-1β release of LPS-primed PBMCs was quantified by HTRF after 1 hr treatment with indicated compound concentrations. IC:50 values were calculated for each compound as indicated. (B) HTRF result after compound permeability test in microvessels. Supernatants were collected after 300 min incubation from basolateral wells. Supernatants were used to treat LPS-primed PBMCs and the induced reduction of IL-1β release was quantified by HTRF. Each compound was tested in duplicate (n=2). Error bars indicate the SD of HTRF measured in quadruplicates. Statistical significance was calculated by using the unpaired T-test.

## Methods

### Cell cultures and conditioned medium

IMR90-4 iPSCs (WiCell) were cultured in mTesR1 (Stemcell Technologies) on Matrigel (Corning) according to WiCell protocols. BMEC differentiation of iPSC cultures based on previously described protocols (Stebbins et al., 2016). For differentiation, 1×10^5^ IPSCs/well were seeded with mTesR1 supplemented with 10 µM Rock Inhibitor Y27632 (Tocris) on 6-well plates (Costar) prior coated with Matrigel. After 3 days of expansion in mTesR1 (D-3 – D-1), cells were cultured in unconditioned medium (UM) for 6 days (D0 – D5). UM contained DMEM/F12-Hepes supplemented with Glutamax, knockout serum replacer, nonessential amino acids, β-Mercaptoethanol (Life Technologies). On D6, endothelial cells were expanded for 2 days by applying endothelial cell medium (EC+/+) based on human endothelial SFM (Life Technologies) supplemented with 1% platelet-poor plasma derived bovine serum (Alfa Aesar), 10 µM all-trans retinoic acid (RA) (Sigma) and 20 ng/ml bFGF (R&D systems). On D8, heterogenous cultures were subcultured with Stempro Accutase (Life Technologies). Primary human brain vascular pericytes (ScienCell™) and primary human astrocytes (ScienCell™) were maintained until passage 3 in either ScienCell™ pericyte medium (PM) or astrocyte medium (AM), respectively according to manufacturer protocols. For the production of conditioned medium, 4×10^6^ HBVP were cultured in 25 cm^2^ PLL coated flasks (Nunc) with either 10 ml endothelial cell medium without RA and bFGF (EC−/−) or ScienCell™ PM. Medium was exchanged after 2 days. First conditioned medium was harvested on day 3 after seeding. Fresh medium was added to cells and harvested again on day 4 after seeding. Conditioned medium was stored at −20°C. To produce conditioned astrocyte EC−/− medium, the same procedure was applied with 1×10^6^ HA per 25 cm^2^ flasks.

### Transwell culture and TEER

On D8, 12-well Transwell filter were collagen/fibronectin coated for 4 hrs before cell seeding. For coating, 4 parts of 1 mg/ml collagen IV (Sigma) were mixed with 1 part of 1 mg/ml fibronectin (Sigma) and 5 parts H_2_O. After coating, 1×10^6^ BMECs were seeded on Transwell filters with EC+/+ by having a final volume of 500 µl in apical and 1500 µl in basolateral chambers. On D9, medium was exchanged to endothelial cell medium without RA and bFGF (EC−/−). For conditioned medium applications, 1500 µl conditioned EC−/− or PM were applied on basolateral sides. For co-cultures, 1×10^6^ HBVP or HA were pre-cultured for 24 hrs on PLL coated basolateral chambers in absence of Transwell filters. Transwell filters with seeded BMEC layers were combined with basolateral co-cultures on D9. TEER between apical and basolateral chambers was measured by using EVOM2 Volt/Ohm meter combined with STX2 electrode set (WPI). Samples were prepared in duplicates and average values were calculated. To avoid temperature effects, all measurements were performed on a 37°C warmed heat plate (Labotect). Blank values, were subtracted from raw data and measurements were corrected for used Transwell filter area.

### Tracer permeability in Transwells

NaF permeability (PE) was assessed at D10 with BMEC layers cultured on 12-well Transwell filters. Medium was exchanged on apical and basolateral chambers prior to assay start. 500 µl EC−/− supplemented with 10 µM NaF (Sigma) was applied on apical chambers. After 15, 30, 45 and 60 min, 150 µl basolateral medium was collected and exchanged with fresh 150 µl EC−/−. After 60 min, 150 µl apical medium was also collected. NaF intensities were measured on SpectraMax Paradigm plate reader (Molecular Devices) equipped with 485/20 nm excitation and 535/25 nm emission filter cartridge (part number 0200-7003). Intensity values were corrected for background and signal loss according to bottom medium dilution. PE was calculated as previously described (Stebbins et al., 2016). Clearance volume = (V_B_*(S_B,t_))/(S_T,60min_), where V_B_ is the volume of bottom chamber (1500 µl); S_B,t_ is the corrected signal of bottom chamber at time, t and S_T,60min_ is the signal of top chamber at 60 min. The linear slope of clearance volume against time was calculated by linear regression for culture (m_c_) and blank filters (m_f_) and PE was calculated as follows: 1/PE=1/m_c_-1/m_f_ (Perriere et al., 2005). PE (cm/min) = [(1/(1/PE))/1000]/area.

### Microvessel culture

Three-lane OrganoPlates® (Mimetas; 4003-400-B) were prepared according a further modified protocol based on the manufacturer’s instructions (Mimetas.com). On three-lane OrganoPlates®, 40 chambers are integrated in a standardized 384-well plate design. Each chamber contains three connected channels (top, gel and bottom) and two PhaseGuides® between top/gel and gel/bottom channels. Top and bottom channels have 320 µm x 220 µm and gel channels 360 µm x 220 µm (w x h) dimensions. PhaseGuides® have a 100 µm x 55 µm (w x h) dimension. For microvessel production, ECM gel was prepared with 3.7 mg/ml NaHCO_3_ (Sigma), 100 mM Hepes (Life Technologies) and 4 mg/ml Collagen-I (Cultrex 3D rat tail collagen-I, R&D systems) on ice. 1.3 µl of ECM gel was pipetted into gel channels. BMECs were harvested at D8 from 6-well plates and 2 µl cell suspension with a concentration between 3.8 − 4.2 × 10^7^ cells/ml was pipetted into top channels. Both top wells and gel wells were filled with 50 µl EC+/+ and plates were incubated at 70° on side for 2 hrs at 37°C and 5 % CO_2_. Plates were further incubated horizontal over-night. Perfusion with 8 min intervals and 7° was started at D9 to apply shear stress inside chamber channels. EC+/+ was exchanged by EC−/− supplemented with HBVP conditioned PM in a ratio of 1:4. Right-bottom wells were filled with 50 µl of the same medium to allow bottom channel filling by capillary force. After visual inspection of bottom channel filling, left-bottom wells were also filled with 50 µl medium. Perfusion was continued for 24 hrs.

### Microvessel TEER

For the TEER measurement of microvessels in the three-lane OrganoPlate®, we adapted the voltage divider method. A standard reference resistor *R* = 490 *k*Ω (± 5% tolerance) was used in series with the microvessels unknown resistance “*r*” to divide a fix reference voltage *V* = 0.4 volt (IC: LT6650) (Supplementary Figure S2E). Low reference voltage allows keeping the current very low through the microvessels. The voltage across the microvessels (V_1_), is probed with a pair of Ag/AgCl electrodes and is amplified (~20 fold) with high input impedance, DC differential amplifier and measured as readout V_0_ (Supplementary Figure S2F). High Input impedance assures that the resistance is measured with high accuracy. Measuring the ohmic resistance of a microvessel is described as follows: From Ohm’s law: If *i* is the current through the resistors (*R and r*) in series then *V* = *i* * (*R* + *r*), and 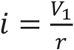. Thus, 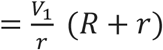, this gives *r* in terms of known parameters *R, V and V*_1_ as: 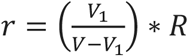. For the differential amplifier when 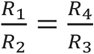 then the readout voltage 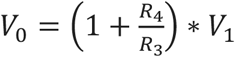. For our chosen resistors *V*_0_ = (1 + 19.69) * *V*_1_ or *V*_1_ = *V*_0_/20.69. Thus 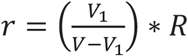 and 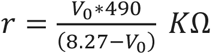.

The cell medium was changed in the chamber before each measurement and the temperature as well as the exposed area of the electrodes were kept constant since the resistance of the thin Ag/AgCl electrode (*𝜙* = 200 *μm*) is sensitive to the surface area of the electrode, temperature and pH. Both Ag/AgCl electrodes were identical and coated properly (measured junction potential difference was ~ 2mV which is within error limit of the voltmeter used). Self-resistance of the electrode (= 136 kΩ) was determined by dipping the electrodes in culture medium with separation of ~1 mm. Ohmic resistance of chambers without microvessel was measured in parallel to the resistance of the chambers containing an intact microvessel. Effective ohmic resistances of microvessel were calculated by subtracting the self-resistance of the electrode and the blank ohmic resistance (chamber without cells). Reported TEER values in Ωxcm^2^ units were calculated by multiplying the effective ohmic resistance of the chamber by the gel-microvessel contact area. Device was calibrated with known standard carbon film resistors (in the place of a chamber containing a microvessel) and the values of resistances matched very closely with the calculated values of the same resistors by the device (Supplementary Figure S2G). The range of obtained TEER values strongly depends on the chamber’s architecture and does not allow a direct comparison to Transwell experiments.

### Microvessel integrity and compound permeability assay

Microvessel integrity was determined on D10 cultures. Top wells were filled with 80 µl HBVP conditioned PM supplemented with 25 % EC−/−, 10 µM NaF, 30 µg/ml 10 kDa DEX-A647. For compound permeability tests, 40 µM compounds were supplemented on top wells. Gel and bottom wells were filled with 20 µl media mix without fluorescent tracer and compounds. Plates were incubated on a perfusion rocker at 37°C and 5% CO_2_. Imaging was performed with a confocal spinning disc CellVoyager 8000 HCA screening microscope (Yokogawa) at 37°C and 5% CO_2_. Acquisition was done with a 10x objective (Yokogawa) within the microvessel lumen (offset from bottom z = 50 µm) with a 488 nm (NaF) and 640 nm (DEX-A647) laser excitation and BP 525/50 and BP 708/75 nm emission, respectively. Both channels were setup with a 50 µm pinhole and 2×2 binning. Automated image analysis was performed by applying a customized ImageJ macro. In brief, images were registered to correct for x, y shifts and background values of both fluorescence channels were subtracted. Average intensity values of both tracers within the top and gel channels were measured by using standardized ROIs. Tracer intensity ratios [gel/top] were calculated. The following compounds were used in this study: CRID3 sodium salt (MCC950 sodium; Cayman Chemicals No.17510; 426.5 MW), Emricasan (SelleckChem S7775; 569.5 MW), Ac-YVAD-CMK (caspase-1 inhibitor II; Cayman chemicals No.10014; 541 MW), Z-DEVD-FMK (SelleckChem S7312; 668.7 MW), Z-VAD(OH)-FMK (caspase inhibitor VI; SelleckChem S8102; 453.5 MW) and Staurosporine (STS; Sigma S5921; 466.5 MW).

### Cluster analysis

K-means cluster analysis was performed by using 6 independent ratio combinations extracted from NaF and DEX-A647 image data at two incubation times: Ratio (R_1_) = gel_Dex_/top_Dex_ (150 min), R_2_ = gel_Dex_/top_Dex_ (300 min), R_3_ = gel_NaF_/top_NaF_ (150 min), R_4_ = gel_NaF_/top_NaF_ (300 min), R_5_ = gel_Dex_/top_NaF_ (150 min), R_6_ = gel_Dex_/top_NaF_ (300 min). To identify the boundaries of leak-tight versus leaky microvessels in the data set, 2 clusters were considered for the analysis. A set of 26 microvessels including DEX-A647 leaky and leak-tight vessels as well as 10 control chambers without cells (no vessel) were finally clustered by using JMP® (2016 SAS Institute Inc.) software. The applied cluster method reached a cubic clustering criterion (CCC) of 2.277.

### HTRF IL-1β quantification

PBMCs (Stem Cell Technologies) were seeded in 384 well plates (Greiner) with a density of 6×10^4^ cells / well. PBMCs were cultured in 30 µl RPMI 1640 (Biochrom) supplemented with L-Alanyl-Glutamine (Biochrom), Hepes (Life Technologies), Ciprofloxacin Kabi solution (Fresenius Kabi), heat inactivated FBS (Sigma/Merck). Cells were primed with a final concentration of 20 ng/ml LPS-EB ultrapure (InvivoGen) for 2 hrs in a cell incubator. 20 µl basolateral supernatants from microvessel permeability tests were added into wells and incubated for 1 hr. Finally, cells were activated for 30 min with a final concentration of 2 mM ATP (Sigma) to trigger IL-1β release. Supernatants were collected to quantify the amount of IL-1β by HTRF according to the manufacturer’s protocol for the human IL-1β detection kit (Cisbio). In brief, the intensities of two different anti-IL1β antibodies labeled with FRET-donor SA-Europium Cryptate and FRET-acceptor XL665 were measured on a SpectraMax Paradigm plate reader (Molecular Devices). The reader was equipped with a single excitation at 340 nm and two emission channels at 616/10 and 665/10 nm (HTRF detection cartridge part number 0200-7011). The instrument optics was set up to 35 pulses with a pulse length of 0.05 ms, pulse delay of 7.7 ms and a 0.2 ms integration time. Standard calibration curves were included according to Cisbio kit. The IL-1β concentration for each tested supernatant was determined according to manufacturer’s data analysis tools (Cisbio). In dose response experiments, monoclonal anti-interleukin-1 beta antibody (R&D systems) was used.

### Immunocytochemistry

Microvessels were fixed with 4 % PFA (Sigma) in PBS (Life Technologies) for 30 min and blocked for 1hr. Blocking solution contained PBS supplemented with 10 % goat-serum (biowest) and 0.1 % Triton-X100 (Sigma). Primary antibodies were incubated for 2 hrs and secondary antibodies for 1 hr. All incubation steps were performed at RT on a perfusion rocker (Mimetas) switching between +7° and −7° every 5 min. 2D BMEC cultures on 96 well plates (Greiner) were fixed with 4 % PFA/PBS for 20 min and blocked with blocking solution for 1 hr. Primary and secondary antibodies were incubated for 1 hr each. All steps were performed at RT. The following antibodies were used: Anti-ZO-1 (Cell Signaling), anti-VE-cadherin (Millipore), anti-Occludin-A488 conjugated (Life Technologies), anti-Claudin-5 (Abcam), anti-Glut1 (Life Technologies), anti-BCRP (Millipore), anti-P-gp (Cell Signaling), anti-Pecam-1 CD31 (Neomarker), goat anti-rabbit Alexa 488 (Life Technologies), goat anti-rabbit Alexa 647 (Life Technologies), goat anti-mouse Alexa 488 (Life Technologies), goat anti-mouse Alexa 647 (Life Technologies). Hoechst (Life Technologies), DRAQ5 (biostatus) or CellMask™ deep red stain (Life Technologies) was used according to manufacturer’s protocols. Imaging including three-lane OrganoPlates® was performed on a confocal spinning disc CellVoyager 8000 HCA screening microscope (Yokogawa). Three excitation channels (405, 488, 640 nm laser) and three emission channels (BP 445/45, BP 525/50, BP708/75) were used. 3D imaging of three-lane OrganoPlates® was performed by using 20x or 40x air objectives (Yokogawa) over a z-range of 210 µm with slicing intervals of 1-3 µm (pinhole 50 µm). Z-stacks were stitched in 5x, 2y (20x objective) for complete 3D microvessel representation by using IMARIS software package (Oxford Instruments). Z-stacks acquired with 40x were stitched in 9x, 3y for completed vessel representation and animation.

## Discussion

In the present work, we describe the development of a human BBB *in vitro* model that combines relevant benchmarks such as brain endothelial protein expression, 3D microvascular structure, wall shear stress, physiological barrier tightness, restricted tracer permeability and cell polarization in a standardized high-content assay format suitable for compound screenings.

The high standard of recent BBB *in vitro* models that provide almost physiological BBB properties is strongly dependent on BMEC supporting cell types such as astrocytes, pericytes and neurons which needs to be co-cultured to generate an artificial microenvironment (Lee et al., 2020; Park et al., 2019; Vatine et al., 2019). However, these procedures require either the parallel maintenance of primary cell types or elaborate cell differentiation in case supporter cell types can be derived from iPSCs (Canfield et al., 2017; Kusuma et al., 2013; Lippmann et al., 2014). In general, co-culture systems can introduce a higher complexity and variance to the assay that could hamper standardized high-content screening applications. Furthermore, the more common procedure of using primary cell sources is often compromised by the limited passage number of the primary material that reduces the total sample number within each screen. In addition, the introduction of primary cells derived from different human donors is challenging for standardization. In the present work, we established a BMEC differentiation protocol that is based on storable primary HBVP conditioned medium that can be produced in bulk. In this way, conditioned medium would allow a standardized and reproducible production of microvessels on a large scale required for screening applications. In contrast to commonly used heterotypical co-culture systems, conditioned media is strongly suggested because it provides more controllable and defined assay conditions as well as robust data interpretation over multiple microvessel plates. Surprisingly, our results indicated that astrocytes are unsuitable for the production of conditioned medium and further studies are required to untangle the molecular mechanisms behind the BBB supporting microvascular environment.

Recent efforts in tissue engineering have led to the development of singular cylindrical endothelial microvessels that can be integrated into a flow system to mimic vascular shear stress (Faley et al., 2019; Katt et al., 2018; Linville et al., 2019; Vatine et al., 2019). These elaborated platforms were able to recreate anatomical, physiological and mechanical forces that endothelial cells experience *in vivo* (Workman and Svendsen, 2020). The drawback of these models is that they are in generally limited in upscaling. In the presented work, we demonstrated the parallel production of 40 perfused microvessels on a single plate in a standardized procedure. The applied bidirectional perfusion by gravity flow generated a shear stress of approx. 1.2 dyne/cm^2^ (Wevers et al., 2018), that is on the lower edge but still within the range of shear stress determined for post-capillary venues (1-4 dyne/cm^2^) in mice (DeStefano et al., 2018). In general, a successful microvessel formation is challenging due to the technical issue of cell clogging within the microfluidic chamber. To this end, applied perfusion 12 hrs after cell seeding was mandatory for successful microvessel formation and unwanted occlusion (data not shown).

The diameter of produced microvessels was approx. 388 µm that was dictated by the chamber design. In general, the production of smaller microvessels to mimic human brain capillaries with a typical diameter of 8-10 µm remains one of the major hurdles in tissue engineering (Jamieson et al., 2017). To successfully reduce the dictated microvessel diameter by the given plate design and to circumvent possible cell clogging effects, promising approaches with self-assembled vascular networks that are based on vasculogenesis or angiogenesis have been reported (Bang et al., 2017; Bogorad et al., 2017; Campisi et al., 2018). To this end, upscaling of such promising *in vitro* models would be in focus of future research.

Recently, organ-chip technology with human iPSC-derived microvessel cultures combined with astrocytes, pericytes and neurons provided physiological relevant TEER values (>1000 W/cm^2^) in a perfused setup (Vatine et al., 2019). The authors applied static, 0.01, 0.5 and 2.4 dyn/cm^2^ and RNA sequencing analysis revealed that expression of tight-junction proteins as well as other endothelial marker were flow dependent, but do not differ between 0.5 and 2.4 dyn/cm^2^ (Vatine et al., 2019). Here we demonstrated that iPSC-derived microvessels in microfluidic plates reached maximal TEER values in the same physiological range by applying 1.2 dyn/cm^2^. Therefore, our produced microvessels provide benchmark barrier properties in a standardized and scalable format by using conditioned media. Moreover, we obtained a linear correlation between microvessel TEER and fluorescence intensity ratios in case of NaF. Due to its molecular properties, NaF has been suggested as the tracer of choice that enables the detection of small changes in BBB tightness associated to small microvascular disruptions *in vivo* (Kaya and Ahishali, 2011). In our study, TEER and NaF intensity data provided a precise and sensitive measure enabling the detection of small changes in barrier tightness of intact microvessels. To our knowledge, this is the first study demonstrating that a quantitative imaging approach is able to recapitulate a TEER measurement in an *in vitro* system.

In general, the detection of barrier disruption in real time is challenging by using fluorescent tracer due to the fact that these dyes need to accumulate in a sufficient way on the basolateral side to detect them by live-cell imaging. In contrast, TEER measurements would allow the detection of rapid barrier disruption or barrier wound healing events in real time by continuous recording. However, in contrast to TEER measurements, image techniques could provide spatial information about the location of vessel damage or wound healing events. In this context, imaging based assays have the major advantage that tracer integrity assays could be combined with additional readouts. Applications to investigate endothelial cell proliferation, wound healing, cell movement, angiogenesis, events of cell death and dynamics of cell-to-cell contact could be included by using appropriate live-cell marker. Furthermore, microvessel imaging with subcellular resolution provides an avenue to investigate cellular transport and could potentially support the development of recent drug-delivery technologies. To this end, we demonstrated BMEC polarization in produced microvessels by addressing the localization of BCRP and Glut-1 highlighting a physiological localization for both proteins as previously described (Iorio et al., 2016; Simpson et al., 2007). Substrates and inhibitors of BCRP include a wide range of various clinically important drugs, and BCRP has been recognized by the U.S. Food and Drug Administration (FDA) to be one of the key drug transporters involved in clinically relevant drug disposition (Mao and Unadkat, 2015). Therefore, we strongly recommend the presented microvessel platform as screening tool to identify BCRP substrates and inhibitors in future.

To evaluate our BBB model for drug permeability tests, we have chosen a collection of caspase inhibitors used in research, but also inhibitors reported in clinical trials. Our permeability analysis included also the broad spectrum peptidomimetic caspase inhibitor Emricasan (IDN-6556, Conatus Pharmaceuticals / Novartis) which has been announced as well tolerated molecule in several clinical trials for liver diseases and liver damage (Kudelova et al., 2015). Notably, Emricasan was previously identified as neuroprotective drug *in vitro* (Xu et al., 2016) and showed promising effects in an ischemia / reperfusion injury rat model (Tian et al., 2018). However, our data showed no permeation of Emricasan through microvessels suggesting it as systematic drug with limited but promising CNS applications to treat stroke related injury in cases were the BBB itself is disrupted (Khatri et al., 2012). CRID3 (MCC950, Inflazome/Roche) that is known as one of the most potent and selective inhibitor of the NLRP3 inflammasome (Coll et al., 2015) was included as BBB permeating reference compound. In line with our results from microvessel permeability tests, it has been demonstrated that CRID3 is known to penetrate effectively the BBB after oral administration in mice (Chen et al., 2017). Moreover, CRID3 treatments in mice demonstrated rescued dopaminergic degeneration in a Parkinsons diseases model (Gordon et al., 2018) and reduction of Aβ accumulation in an Alzheimer’s disease model (Dempsey et al., 2017). During last years, several anti-inflammatory compounds against a wide range of CNS and non-CNS diseases were announced for clinical trials including molecules inspired by the structure of CRID3, highlighting the potential of this class of molecules. Besides CRID3, we have identified two additional and so far, unpublished anti-inflammatory compounds (presented as cmpd-A and -B) that were able to penetrate produced microvessels and further studies will be required to evaluate these molecules.

In summary, our BBB *in vitro* platform based on the differentiation of human iPSC-derived BMECs by applying conditioned media without compromising the physiological barrier properties. Thus, the here presented platform enables improved throughput for compound screening to finally better predict BBB drug-permeability during lead optimization. The here used microfluidic plates contain 40 single chambers per plate and microvessels could be fabricated by automated liquid handling systems that would allow cost effective and standardized workflows required for large screenings. Every chamber on the plate is addressable for high-content imaging applications providing the additional option of live-cell acquisition and quantitative microscopy which could also support the development of novel drug transport or carrier systems.

In general, the combination of our microvessel system with a cellular readout may prove useful to discriminate between BBB permeable and non-permeable anti-inflammatory compounds in existing compound libraries. To this end, the identification of BBB permeable candidates in particular could potentially accelerate the drug development for neurodegenerative diseases by trying to repurpose already approved drugs. Moreover, implementation of microvessels from specific patient derived iPSCs could be used to model various CNS diseases that comprise the BBB function. Furthermore, differences in patient responses to drugs could be simulated *in vitro* as well as personalized medicine applications can be developed in the same context.

## Acknowledgments

We thank Ritah Gwokyalya for her technical assistance. We thank Mansoureh Aghabeig and Christoph Moehl for their support in data analysis. We also thank Josephine Blersch, Dominik Stappert and Hardik Doshi for critical reading of the manuscript.

## Author Contributions

SF and PD coordinated the study, designed the experiments and wrote the manuscript. SF, BK and WR performed the experiments and analyzed the data. SKK performed data analysis and developed the TEER device. PD proposed the idea and supervised the project. EF provided general project coordination and intellectual input. All authors revised the manuscript and approved the submitted version.

## Supplementary information

**Supplementary Figure S1.**
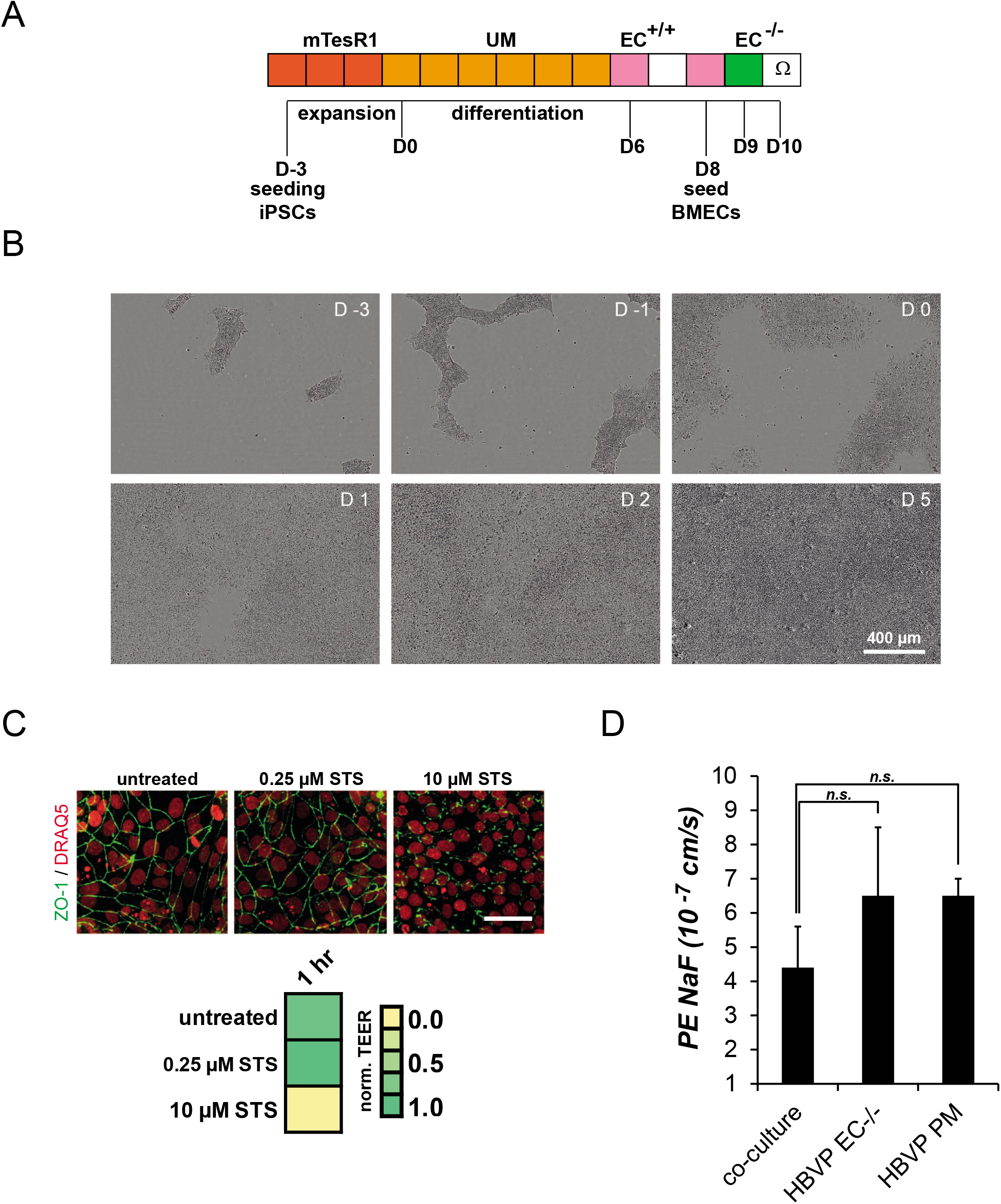
(A) BMEC differentiation protocol. iPSCs were seeded as singularized cells onto Matrigel coated 6-well plates and expanded for 3 days in mTesR1 medium. On D0 medium was changed to unconditioned medium (UM) resulting in a mixed endothelial cell / neuronal progenitor cell population. On D6, endothelial cells were selectively expanded by changing to endothelial cell medium supplemented with RA and bFGF (EC+/+). To purify BMECs, heterogeneous cultures are subcultured at D8 onto collagen/fibronectin-coated Transwell filters. On D9, EC+/+ was replaced by endothelial cell medium without RA and bFGF (EC−/−). (B) Representative brightfield images of differentiating cells from D-3 to D5. Scale bar equals 400 µm. (C) Normalized TEER measured on Transwell filters and corresponding immunocytochemistry of STS treated BMEC cultures. Cells were stained for nuclei shown red (DRAQ5™) and tight-junctions shown in green (anti-ZO-1). Scale bar equals 50 µm. (D) NaF permeability (PE) determined for different BMEC Transwell cultures. BMECs were either cultured with HBVP conditioned medias (EC−/− or PM) or co-cultured with basolateral HBVPs. Mean values and standard deviation from triplicates are shown. Statistical significance was calculated using the unpaired T-test (n.s. p>0.05).

**Supplementary Figure S2.**
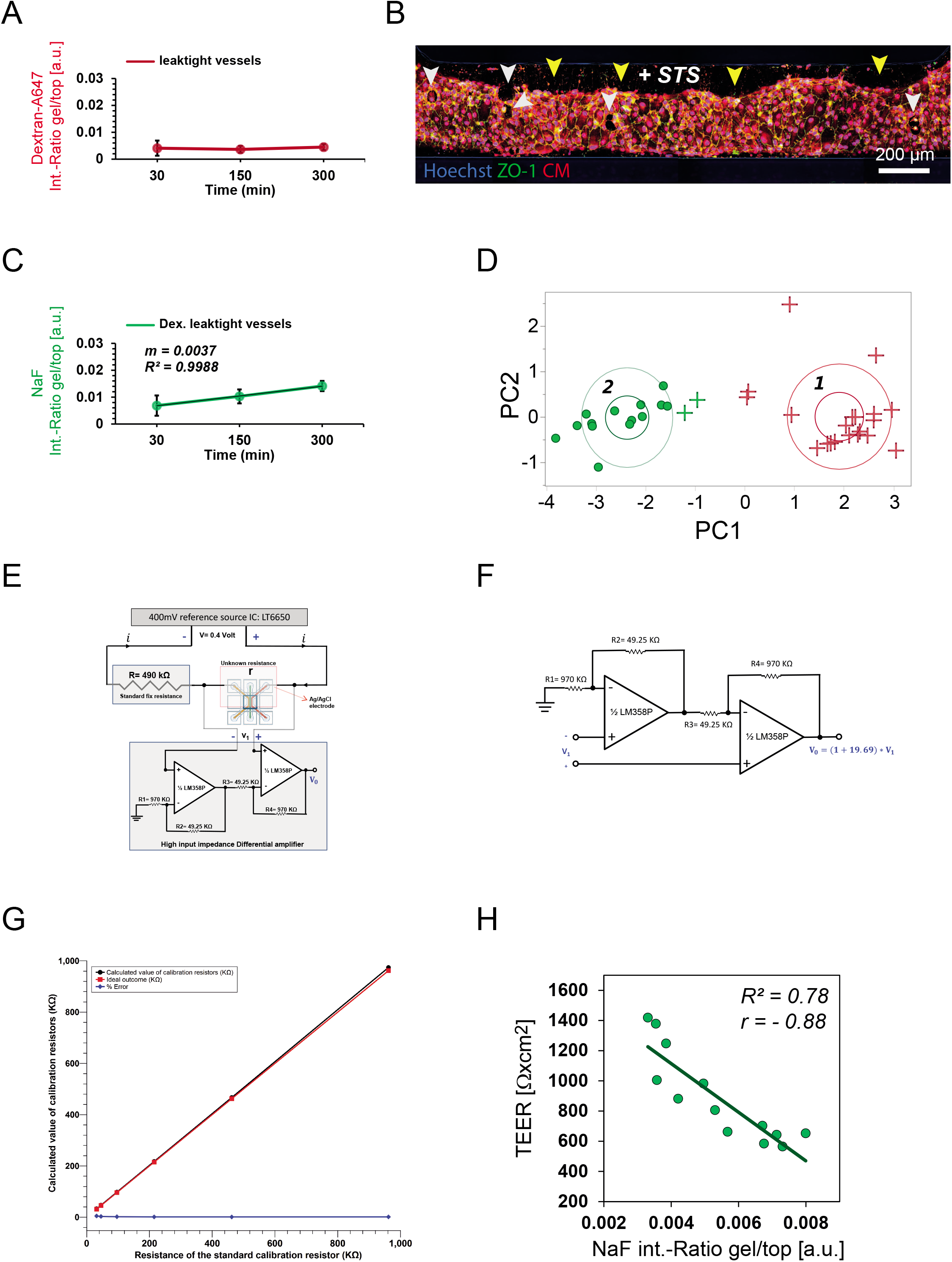
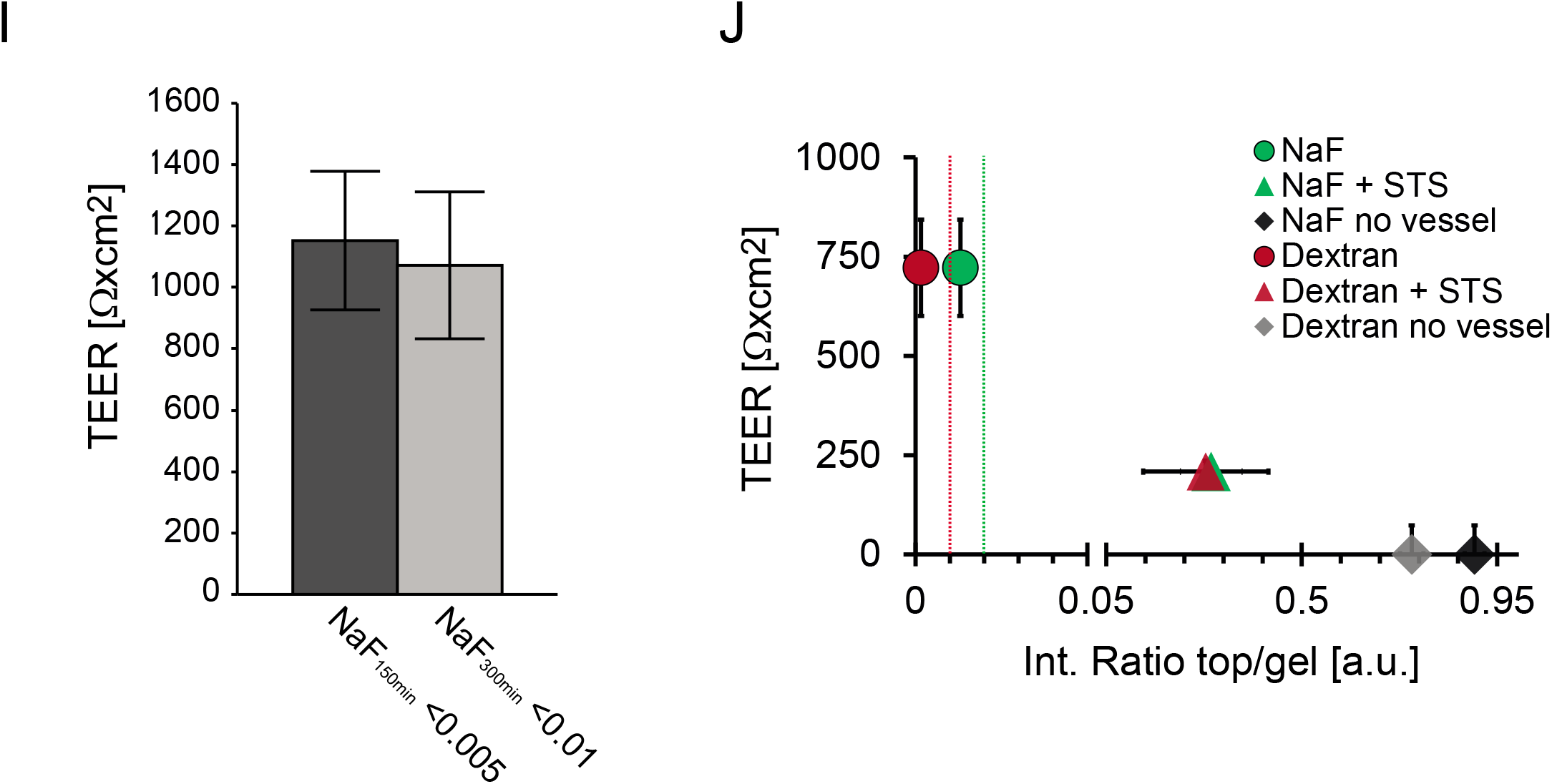
(A) DEX-A647 mean intensity ratios (gel/top) over time with a different y-scale as depicted in (Figure 3B). DEX-A647 leak-tight microvessels with a constant ratio over time (< 0.01) are shown (red line). Error bars indicate the SD (n=6). (B) Microvessel treated by 20 µM STS for 120 min were fixated by 4% PFA/PBS and stained by immunocytochemistry for nuclei (Hoechst), tight-junction marker (ZO-1) and cytoplasm marker (CellMask™). Loss of microvessel adhesion (yellow arrowheads) and vessel holes (white arrowheads) induced by STS are indicated. (C) NaF mean intensity ratios (gel/top) over time with a different y-scale as depicted in (Figure 3C). NaF intensity ratios of preselected DEX-A647 leak-tight microvessels are shown (green line). Error bars indicate the SD (n=6). Slope (*m*) and R^2^ from linear regression of data points are shown. (D) K-means cluster analysis by using tracer intensity data with 6 independent variables as described in the methods section. 2D scatterplot of principal component 1 (PC1) versus PC2 is indicated. Cluster members of cluster 1 are shown in red and members of cluster 2 in green. Leak-tight microvessels are shown as green dots which were defined after excluding those cluster members found at the outer boundaries of cluster 2 (fraction with a green cross). Leaky microvessels are indicated by a red or green cross. A total of n=36 microvessels were analyzed including n=10 control chambers without microvessel (no vessel). Chambers without a microvessel clustered without any exception into cluster 1 (leaky). From a total of 26 microvessels, 14 candidates were finally defined as leak-tight. (E-F) Layout and circuit of the customized microvessel TEER device. (G) Calibration and testing of the TEER device with known standard resistors. (H) Correlation of TEER versus NaF ratios measured in leak-tight microvessels after 150 min incubation. Green line indicates the linear regression of NaF/TEER data points. R^2^ of the regression analysis and the calculated correlation coefficient r is indicated in the graph. (I) Average TEER of leak-tight microvessels calculated according to different NaF thresholds (R_gel/top_ NaF_150min_ < 0.005 and R_gel/top_ NaF_300min_ < 0.01). Error bars indicate the SD of n=6. (J) Scatter plot of average NaF (and DEX-A647) intensity ratios (on x-axis) versus average TEER (on y-axis). Data of leak-tight microvessels before (dots) and after 60 min STS treatment (triangles) are shown in the graph. Data from chambers without microvessel (no vessel) are shown as grey diamonds in comparison. Error bars show the SD of (n=2). Thresholds defined by K-mean clustering analysis were applied to distinguish between leak-tight and leaky microvessels by using NaF and DEX-A647 ratio data. Threshold for NaF (R_gel/top_ NaF_300min_ < 0.02) indicated as green line and thresholds for DEX-A647 (R_gel/top_ DEX_300min_ < 0.01) indicated as red line. By considering the fluorescent tracer data, previously leak-tight microvessels shifted into the leaky cluster after incubation with STS. The result was in line with a measured decrease in TEER after STS.

**Supplementary Figure S3.**
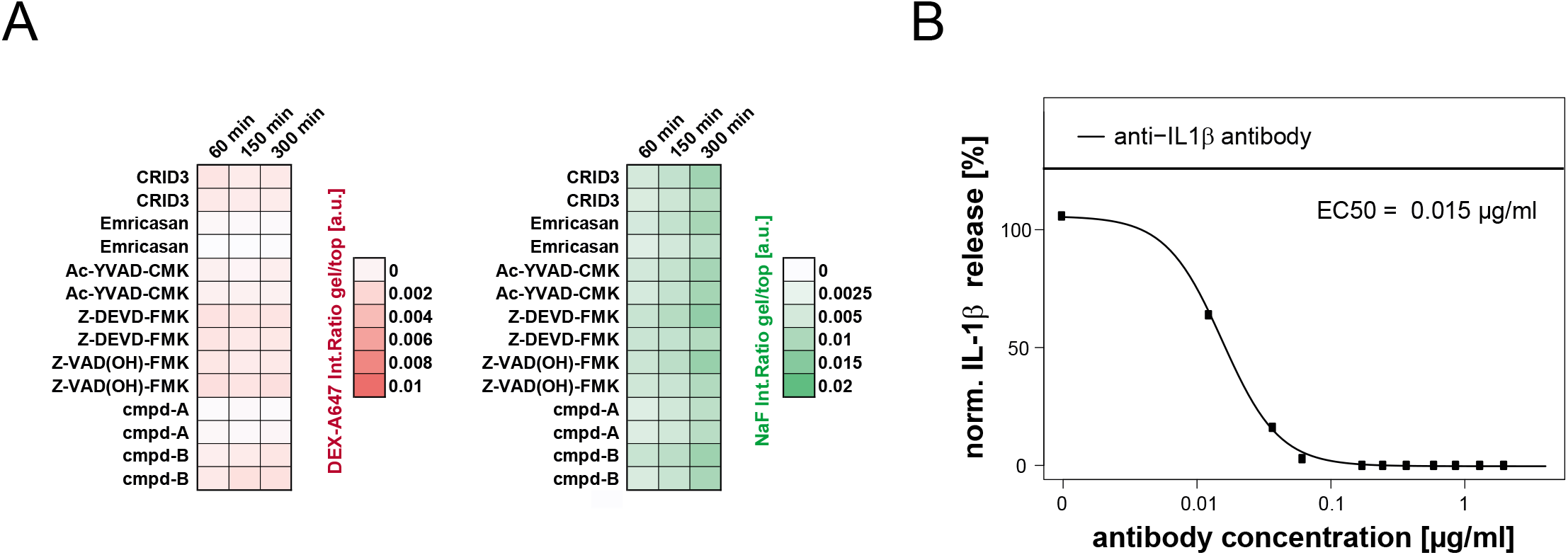
(A) Time profile of DEX-A647 (left) and NaF (right) intensity ratios (top/gel) of microvessels used in the compound permeability assay. Each compound was tested in duplicate. (B) Quantification of IL-1β in PBMC supernatants by HTRF. Titration of anti-IL1β antibody results in a concentration dependent neutralization of released IL-1β.

**Supplementary Video V1**

Animation of representative images recorded during BMEC differentiation from D-3 to D5.

**Supplementary Video V2**

Animation of a 3D reconstructed microvessel from imaging data. The microvessel was stained for nuclei (Hoechst) in blue, tight-junctions (ZO-1) in green and cytoplasm CellMask™ in red.

